# Extracellular RNA induce neutrophil recruitment via endothelial TLR3 during venous thrombosis after vascular injury

**DOI:** 10.1101/2024.01.18.576322

**Authors:** Maria Y. Najem, Ryan N. Rys, Sandrine Laurance, François-René Bertin, Virginie Gourdou-Latyszenok, Lénaïck Gourhant, Lauriane Le Gall, Rozenn Le Corre, Francis Couturaud, Mark D. Blostein, Catherine A. Lemarié

## Abstract

**Background:** Venous thromboembolism is associated with endothelial cell activation that contributes to the inflammation-dependent activation of the coagulation system. Cellular damages are associated with the release of different species of extracellular RNA (eRNA) involved in inflammation and coagulation. TLR3, which recognizes (viral) double-stranded RNA, single-stranded RNA, and also self-RNA fragments might be the receptor of these eRNA during venous thromboembolism. We investigate how eRNA regulate endothelial function through TLR3 and contribute to venous thromboembolism.

**Methods and Results:** Thrombus formation and size in WT and TLR3 deficient (-/-) mice were monitored by ultrasonography after venous thrombosis using the FeCl_3_ and stasis models. Mice were treated with RNase1, poly(I:C) or RNA extracted from murine endothelial cells (eRNA). Gene expression and signaling pathway activation were analyzed in HEK293T cells overexpressing TLR3 in response to eRNA or in HUVECs transfected with a siRNA against TLR3. Plasma clot formation on treated HUVECs was analyzed. Thrombosis exacerbated RNA release in vivo and increased RNA content within the thrombus. RNase1 treatment reduced thrombus size compared to vehicle-treated mice. Poly(I:C) and eRNA treatments increased thrombus size in WT mice, but not in TLR3^-/-^ mice, by bolstering neutrophil recruitment. Mechanistically, TLR3 activation in endothelial cells promotes CXCL5 secretion and neutrophil recruitment in vitro. eRNA triggered plasma clot formation. eRNA mediate these effects through TLR3-dependent activation of NFκB.

**Conclusions:** We show that eRNA and TLR3 activation enhance venous thromboembolism through neutrophil recruitment and secretion of CXCL5.

## Introduction

Venous thromboembolism (VTE), which includes deep-vein thrombosis (DVT) and pulmonary embolism (PE), is a common condition affecting 1% to 2% of the population, with an annual incidence of 1 in 500^1^. VTE can lead to death through PE in 10% of cases, result in the post-thrombotic syndrome in 30% of VTE presenting as DVT, characterized by chronic leg pain, swelling, and ulceration, or in chronic pulmonary hypertension in 4% of cases resulting in significant chronic respiratory compromise and mortality^2,3^. VTE is a multifactorial disease involving acquired or inherited risk factors. Among these, surgery and trauma are among the most common risk factors associated with VTE. Both VTE and acute inflammation share an increase in endothelial and platelet activation raising the possibility that factors derived from the inflamed endothelium and/or activated platelets might contribute to the thrombotic response. Moreover, studies showed that innate immunity is involved in the formation of venous thrombosis^4–6^. Interestingly, data obtained in mice suggest that aberrant activation of immunity during thrombosis is a key event in the initiation and propagation of venous thrombosis. Indeed, it is now well established that VTE starts as sterile inflammation with a massive recruitment of neutrophils and monocytes^4,7^.

Sterile inflammation is associated with the release or production of damage-associated molecular pattern (DAMP) including DNA, RNA or ATP. Toll-like receptors (TLRs) are a critical component of the innate immune system. They participate in host defense against foreign pathogens via the pathogen-associated molecular pattern (PAMP) recognition. TLRs respond not only to foreign pathogen structures but also to host-derived molecules that are produced by injured tissues and cells^8^. Studies demonstrated that short RNA sequences similar to those seen in RNA interference or mRNA are sufficient to stimulate TLR3^9^. TLR3 was described as the receptor of double-stranded (ds)RNA, a by-product of viral replication, in the endosome. In addition, TLR3 has been increasingly linked to tissue damage. Hence, endogenous RNA released from healthy and apoptotic cells or damaged tissue has been correlated with TLR3 expression and signaling^9,10^. TLR3 was proposed to be the sensor of endogenous RNA.

Activation of endothelial TLR3 is involved in endothelial dysfunction and disruption of the hemostatic balance in endothelial cells^11,12^. TLR3 deficiency was shown to protect mice against inflammatory responses in both septic and necrotic processes in the damaged gastrointestinal tract^13^. In addition, TLR3 deficiency reduced myocardial infarction damage after ischemia/reperfusion^14^. Reports indicate that extracellular RNA (eRNA), generated during tissue damage and inflammation in the presence or absence of infection, can exert pro-thrombotic and pro-inflammatory properties in the vascular system. eRNA were found to augment the (auto-) activation of proteases in the contact phase pathway of blood coagulation such as factors XII and XI, both exhibiting strong RNA binding^15^. eRNA were also found to mediate leukocyte adhesion *in vivo* and *in vitro* by promoting vascular cell adhesion molecule-1 (VCAM-1) expression from endothelial cells and tumor necrosis factor alpha (TNFα) secretion by monocytes^16^. In addition, eRNA accumulate in atherosclerotic plaques and induce a pro-inflammatory phenotype from macrophages^17^. Of note, pro-inflammatory and pro-thrombotic agents can induce the release of RNase1 from endothelial cell granule stores (Weibel-Palade bodies). RNase1 treatment was shown to reduce atrial thrombosis, infarct size after stroke, as well as limit cerebral edema and atherosclerosis^14,15,17,18^. However, it remains unknown if eRNA can impact endothelial cell functions during VTE. Thus, we hypothesize that eRNA may be involved in the inflammatory development of VTE via TLR3.

## Materials and Methods

### Animals

Eight-to twelve-week-old male C57Bl/6NJ (stock number 005304) and B6N.129S1-Tlr3^tm1Flv^/J (TLR3 deficient, C57Bl/6J strain backcross 10 generation, stock number 009675) were purchased from Jackson Laboratory. Mice were raised in normal cages under SPF conditions. All procedures described were approved by the Animal Care Committee at McGill University, the local animal welfare committee and the French Ministry for Higher Education and Research according to the European Community guidelines. Only mice developing venous thrombosis following the FeCl_3_ or the stasis models were included in the study. Comparisons were done between WT or TLR3^-/-^ mice and according to the treatments after thrombosis induction.

### Treatments and venous thrombosis models

Prior to thrombosis induction, mice received an intravenous injection of either vehicle, SYTO RNA Select (25 μmol/L, Molecular Probes), RNase I (0.3 μg/μL, Fermentas), polyinosine-polycytidylic acid (poly(I:C), 5 μg/mL, Invivogen) or mRNA prepared from endothelial cells from WT mice (10 μg/mL). The ferric chloride (FeCl_3_, 0.18 M) model of thrombosis was used as previously described^19^. Thrombus cross sectional area was measured using high frequency ultrasonography (HFUS; Vevo 770; Fujifilm; VisualSonics) and a 40 MHz mouse probe as previously described^20,21^. In separate groups of mice, thrombi were dissected out directly from the inferior vena cava (IVC), blotted dry and weighted.

Additionally, when indicated, venous thrombosis was induced using the stasis model as previously described^20^. Briefly, the inferior vena cava and all side branches distal to the left renal vein were ligated using 7-0 Prolene suture (Ethicon, New Jersey). Cross sectional area was measured using high frequency ultrasonography (Canon) 48h after the procedure. Mice were treated with vehicle, poly(I:C), RNase I or eRNA as described above.

### Quantification of plasma DNA and RNA

Plasmatic levels of DNA were quantified using Quant-IT OliGreen ssDNA reagent and kit (Thermo Fisher Scientific) according to the manufacturer’s instructions. Briefly, plasma poor platelet was prepared from whole blood by centrifugation at 1, 000 g for 10 minutes at 4°C and an additional 15 minutes at 10, 000 g at 4°C. DNA concentrations in plasma were quantified using a fluorescent stain provided in the kit. DNA concentration was determined from a standard cure generated with oligonucleotide standard provided in the kit, an 18-base M13 sequencing primer.

Plasmatic concentrations of RNA were quantified using Quant-IT RiboGreen RNA reagent and kit (Thermo Fisher Scientific) according to the manufacturer’s instructions. Briefly, circulating RNA was quantified using the ultrasensitive fluorescent RNA stain, RiboGreen. RNA concentrations of samples were determined from a standard curve generated with ribosomal RNA standard provided in the kit.

### Histology

Thrombi were embedded in OCT and serial 7-μm frozen sections were cut using cryostat and transferred onto gelatin-coated slides. For histology, slides were stained by hematoxilin and eosin (H&E). Thrombus area was quantified using ImageJ software (NIH). For each sample, several cross sections were quantified along the longitudinal axe of the thrombus and then averaged.

For experiments using the ligation model, the Carstairs’ method was used to distinguish red blood cells (yellow), fibrin (orange-red) and platelets (grey). Surface area was quantified on several longitudinal sections of Carstairs’ staining using ImageJ software (NIH) and then averaged.

### Immunofluorescence

For immunofluorescence staining, samples were incubated with 0.5% Triton X-100 for 10 min followed by 10% BSA for 30 min at room temperature. The sections were incubated for 1 h at room temperature in 0.3% BSA in PBS with a rat anti-mouse Ly6G (BD Bioscience), a rat anti-MOMA-2 (Abcam) and rabbit anti-mouse CitH3 (Abcam) antibodies. After washing, sections were incubated with the 555- and 488-nm Alexa Fluor-conjugated secondary antibodies in 0.3% BSA in PBS for 1 h at room temperature. Nuclei were labeled with DAPI.

### ELISA

CXCL5/LIX in culture media of WT and TLR3^-/-^ endothelial cells were determined by enzyme-linked immunosorbent assay (ELISA) according to the manufacturers’ instructions (R&D Systems).

For the detection of TAT complex level in platelet-poor plasma, mouse TAT complex ELISA kit (Abcam) was performed according to the manufacturer’s instructions.

### Endothelial cell culture and cell transfection

Endothelial cells from WT and TLR3^-/-^ mice were prepared as previously described^19,22^. Briefly, endothelial cells were purified by two consecutive immuno-selection procedures using magnetic beads conjugated with anti-ICAM-2 antibody (BD Pharmingen) and plated into 75 cm^2^ tissue culture coated flasks. Endothelial cells were grown in endothelial cell basal medium (EBM-2) supplemented with an endothelial cell bullet kit (EGM-2) (Lonza) and used between passage 3 and 5 in tissue culture dishes coated with 0.1% gelatin and maintained at 37°C in a humidified incubator at 5% CO_2_. For experiments, endothelial cells were incubated for 4h with 5 μg/mL poly(I:C) or with 20 μg/mL of extracellular RNA prepared as described below.

HUVECs (Promocell) were grown in the same conditions as murine endothelial cells and used between passage 3 and 6. HUVECs were transfected with a negative control small interference (si)RNA or siRNA (150 pmol) targeting TLR3 (Santa Cruz Biotechnology) and then incubated with 5 μg/mL poly(I:C). A maximum of gene silencing was observed after 48 h of transfection as shown by quantitative real-time PCR. When indicated HUVECs were also treated with 20 μg/mL of extracellular RNA prepared as described below. In a separate set of experiments, HUVECs were transfected with a negative control siRNA and a siRNA targeting TLR3 (5 nM) from IDT for 48h prior to incubation with 5 μg/mL poly(I:C) or 20 μg/mL eRNA extracted from HEK293T. HUVECs were treated for 45 minutes for Western Blot and 4h for real-time qPCR analysis.

### Isolation of human neutrophil

Human neutrophils were isolated from whole blood. Blood was drawn in EDTA tubes. Blood was layered on lympholyte-poly (Cedarlane) and centrifuge 30 min at 500x g. After centrifugation, two leukocyte bands were visible. The polymorphonuclear band (lower band) was collected and resuspend in culture media. Cells were maintained in RPMI 1640 medium supplemented with 10% FBS, 10 mmol/L HEPES, 1.5 mM L-glutamine, 100 units/mL penicillin, and 100 μg/mL streptomycin (Wisent).

### Migration assays

Twenty-four-well plates and 8-μm pore-size polyvinylpyrrolidone free polycarbonate membranes (VWR) were used. 700 μL of conditioned media from HUVECs transfected with siRNA control or against TLR3 (Santa Cruz Biotechnology) and incubated with poly(I:C) was added in the bottom chamber. Neutrophil suspension (1 x10^6^ cells/mL) was added to the top wells. EGM-2 was used as a negative control. The plate was incubated overnight at 37°C. Membranes were then removed, fixed, and stained using DAPI. The number of neutrophils that migrated to the lower side of the membrane was counted in 10 random microscope fields using the x20 objective (Leica). Neutrophils that passed through the membrane were quantified. When indicated migration was analyzed in presence of SB225002, CXCR2 antagonist (1 nM; Tocris).

### HEK 293T plasmid transfection

HEK 293T were transfected with a human TLR3 plasmid or an empty vector (Addgene). Briefly, 400 000 cells were plated in 6 well plates in high glucose DMEM supplemented with 10% FBS. The next day, 2 μg of pcDNA3 or TLR3 plasmids were mixed with the transfection reagent (JetPrime, Ozyme) and added to the cells for 48h. Cells were then treated with 5 μg/mL poly(I:C) or 20 μg/mL eRNA for 30 min for Western Blot and for 4h for real-time qPCR analysis.

### RNA isolation, Reverse-Transcription and quantitative real-time PCR

Gene expression was evaluated in murine endothelial cells, HUVECs and HEK293T by quantitative real-time PCR (qRT-PCR). Total RNA was extracted from cultured cells using a commercial kit following the manufacturer’s instructions (Genaid, Froggabio or Nagel-Macherey). One μg of total RNA was reverse-transcribed as per the manufacturer’s instructions (VWR and BioRad). The SYBRgreen intercalant was used for amplification detection with the Fast SYBRgreen master mix (Applied Biosystems and BioRad). Primers were designed using the Primer Express Software (Applied Biosystems). The TATA box binding protein (TBP) gene for human cells and S16 for murine cells were used for normalization. Fold changes were calculated using the ΔCt method and results were expressed as fold change ± SEM of five to seven independent experiments. Murine and human primer sequences are listed in Table I.

**Online Table I.**
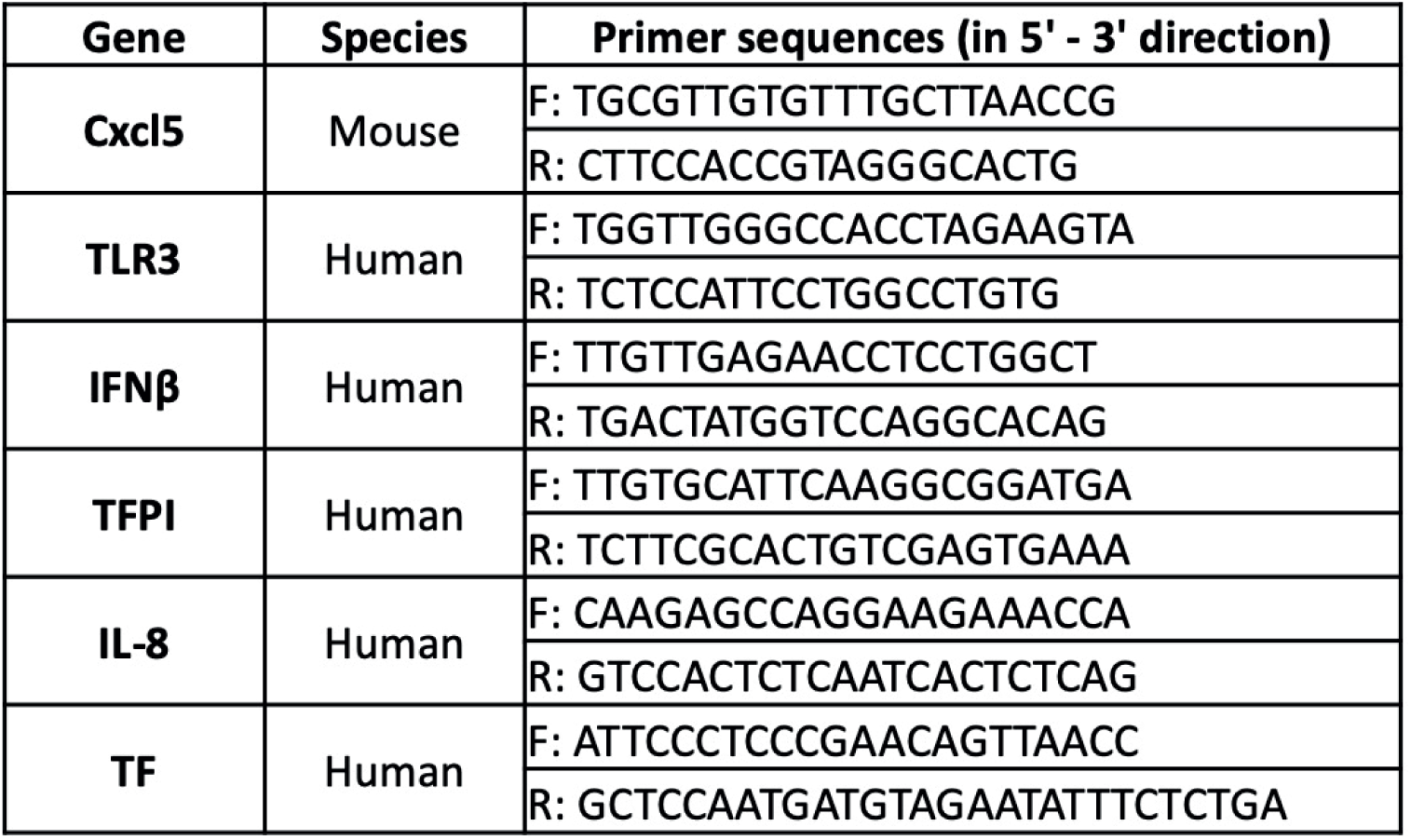
Real time PCR primer sequences.

RNA used to treat HUVECs or HEK293T was isolated from HEK293T using a commercial kit following the manufacturer’s instructions (Nagel-Macherey). RNA used to treat mice and murine endothelial cells was extracted from murine endothelial cells and 3T3 cells, respectively using the same method.

### Western Blot

HUVECs were lysed with 100 µL of whole cell lysis buffer (1% Igepal CA-630, 0.5% deoxycholic acid, 0.1% sodium dodecyl sulfate and 1% Triton X-100) supplemented with 1 M dithiothreitol, 100 mM sodium orthovanadate and complete protease inhibitor cocktail (Roche). HEK293T were lysed with commercially available mammalian protein extraction reagent buffer (Fisher Scientific). Twenty-five µg of protein were separated by 10% sodium dodecyl sulfate polyacrylamide gel electrophoresis (SDS-PAGE) and transferred to nitrocellulose membranes (Fisher Scientific). Western blot analysis was performed with the following antibodies: rabbit anti-phospho-NFκB (1:1000, Cell Signaling), rabbit anti-NFκB (1:2000, Cell Signaling Technology) and mouse anti-GAPDH (1:2000, Santa Cruz Technology). Anti-rabbit coupled to HRP was used as secondary antibodies (1:2000, Dako). Signals were revealed by chemiluminescence (Thermo Fischer) or the GeneGnome molecular chemiluminescence imager system (Syngene) and quantified by densitometry with Quantity one software (Bio-Rad).

### Plasma clot formation

HUVEC were seeded at a density of 100,000 cell per well in 96 well plate. The next day, cells were serum-starved (EBM-2) and stimulated overnight with 1 U/mL thrombin, 5 μg/mL poly(I:C) or 20 μg/mL eRNA. Re-calcified (10 μM) plasma from healthy individuals was added to the wells and formation of fibrin was monitored by measuring OD at 405nm every 2 min for 2h. Curve analyses were done according to the ISTH SSC guidelines^23^.

### Statistical analysis

Data are presented as means ± SEM from multiple independent experiments. Normality was assessed using the d’Agostino-Pearson test. When comparing two experimental groups (WT mice with or without RNase I treatment), Student’s t test was used. When comparing more than 2 groups, with different genotypes (WT or TLR3) and different treatments (vehicle, poly(I:C) or eRNA), two-way ANOVA followed by Tukey’s multiple comparisons test were done. When normality was not present, Kruskal-Wallis test was performed. GraphPad Prism software was used to analyze data and to generate graphs (version 9.0.1). A value of P < 0.05 was considered statistically significant.

## Results

### TLR3 is required for venous thrombosis induced by eRNA

We chose the FeCl_3_ model to induce thrombosis to generates cellular damages, which we hypothesized might exacerbate the release of extracellular RNA from the vascular wall during venous thrombosis. The presence of eRNA in venous thrombus induced by FeCl_3_ was assessed by the injection of SYTO RNASelect. eRNA were detected within the thrombus of WT mice (Figure 1A). To evaluate the role of eRNA in venous thrombosis, WT mice were treated with RNase1 prior to thrombosis induction. Thrombus size was significantly decreased by RNase1 treatment compared to vehicle (Figure 1B). Accumulation of monocytes and neutrophils within thrombi, as shown by MOMA-2 and Ly6G staining, was also reduced by RNase1 treatment (Figure 1C).

**Figure 1.**
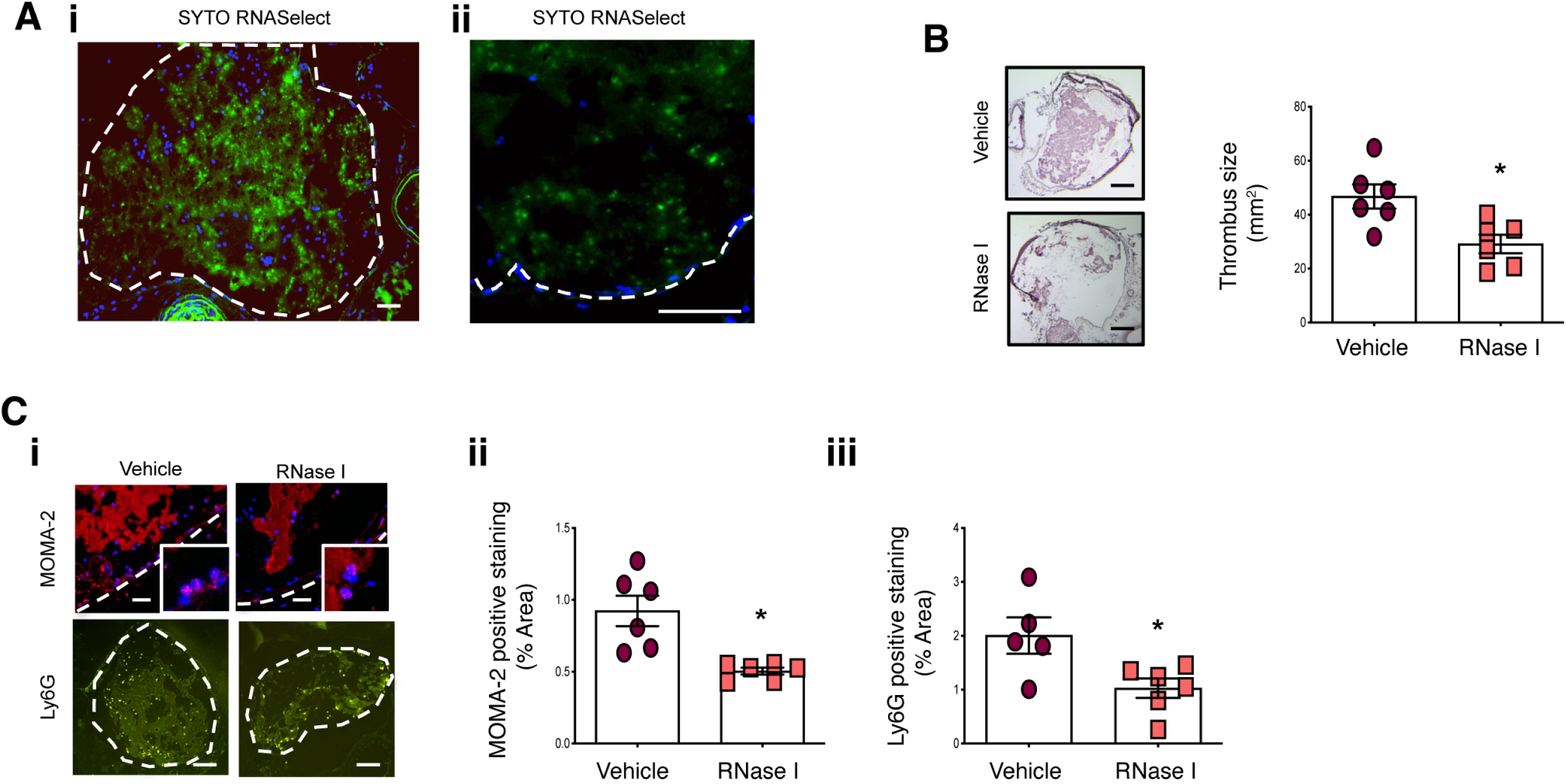
Extracellular RNA promote venous thrombosis. (A) Extracellular RNA (eRNA) is present within thrombi after 30 minutes of vascular injury following FeCl_3_ (i: magnification 10x; Bar=50 μm; ii: magnification x40; Bar=10 μm). (B) RNase I treatment reduces thrombus size in mice after vascular injury. (C) Treatment with RNase I is associated with reduced leukocyte infiltration as shown by MOMA-2 staining for monocyte/macrophage (Bar=5 μm) and Ly6G staining for neutrophils (Bar=50 μm). (N=6) *P<0.05.

Next, we evaluated if direct activation of TLR3 promotes venous thrombosis. Injection of poly(I:C), TLR3 synthetic ligand, increased thrombus size in WT mice (Figure 2A). In addition, poly(I:C) enhanced the number of neutrophils and neutrophil extracellular trap (NET) formation within thrombi of WT mice (Figure 2B, C). Interestingly, these effects were blunted in TLR3^-/-^ deficient mice.

**Figure 2.**
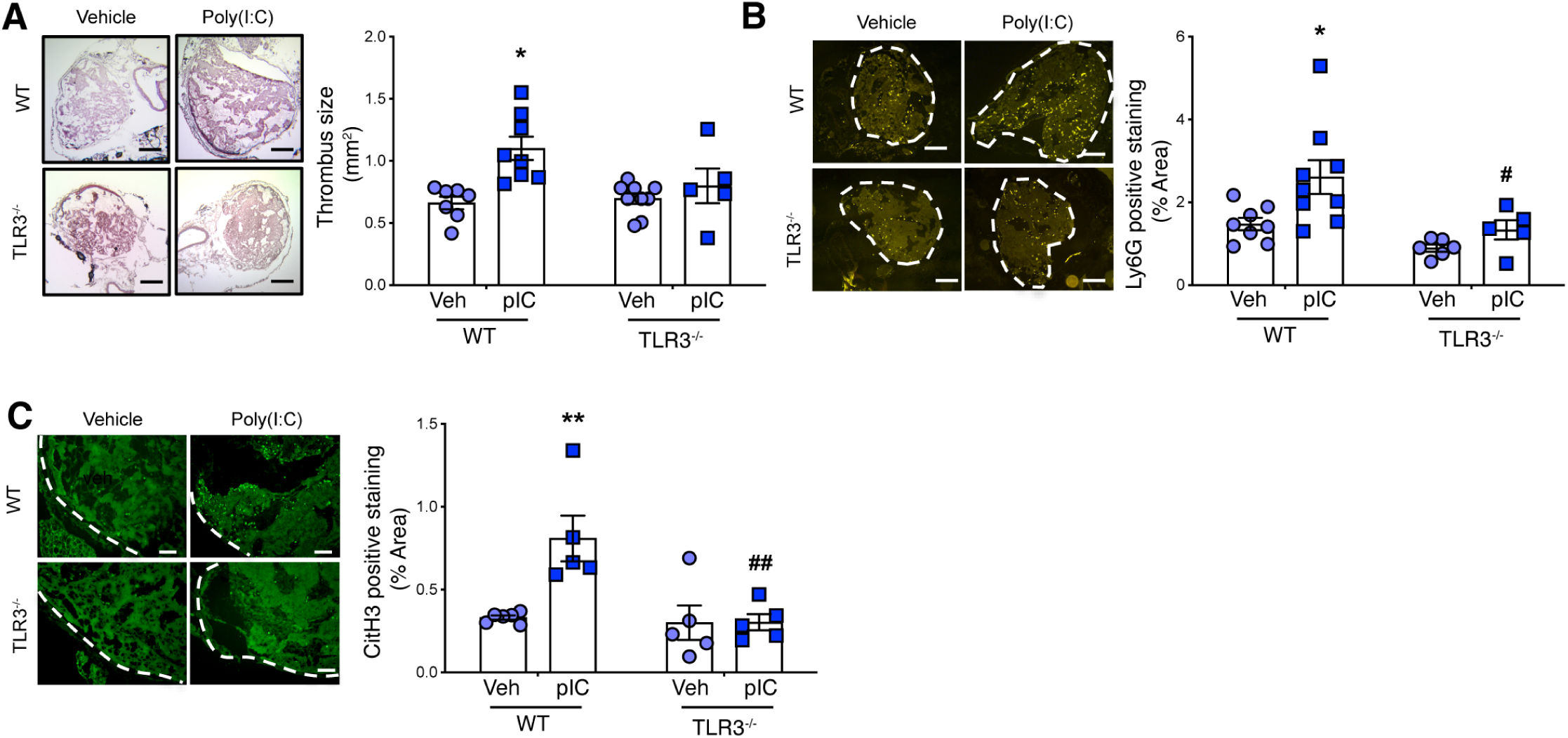
Activation of TLR3 promotes venous thrombosis. (A) Injection of poly(I:C) increased thrombus size in WT but not TLR3^-/-^ mice (magnification 10x; Bar=50 μm; N=5-9). (B) Poly(I:C) treatment increased infiltration of neutrophils within thrombi of WT but not TLR3^-/-^ mice (magnification 10x; Bar=50 μm; N=5-9). (C) Poly(I:C) induced the formation of NETs within thrombi of WT but not TLR3^-/-^ mice (magnification 20x; Bar=5 μm; N=5-6) *P<0.05 or **P<0.01 WT vehicle vs WT poly(I:C), #P<0.05 or ##P<0.01 WT poly(I:C) vs TLR3^-/-^ poly(I:C).

To investigate the role of eRNA, WT and TLR3^-/-^ mice received an injection of 10 μg of eRNA extracted from murine endothelial cells. Plasma concentrations of circulating RNA were increased after induction of thrombosis with FeCl_3_ in mice receiving eRNA injections compared to vehicle-treated animals (Figure 3A). Using high frequency ultrasound, we showed that injection of eRNA significantly increased thrombus size in WT mice. Interestingly, in TLR3^-/-^ mice, treatment with eRNA did not induce further increase of thrombus size compared to vehicle (Figure 3B, C). Thrombus weight quantifications confirmed that injection of eRNA increased thrombus size in WT but not TLR3^-/-^ mice (Figure 3D). In addition, eRNA increased neutrophil recruitment in WT but not TLR3^-/-^ mice (Figure 3E). Finally, TAT complexes quantifications were similar in all treatment groups following FeCl_3_ thrombosis induction (Suppl Figure IA).

**Figure 3.**
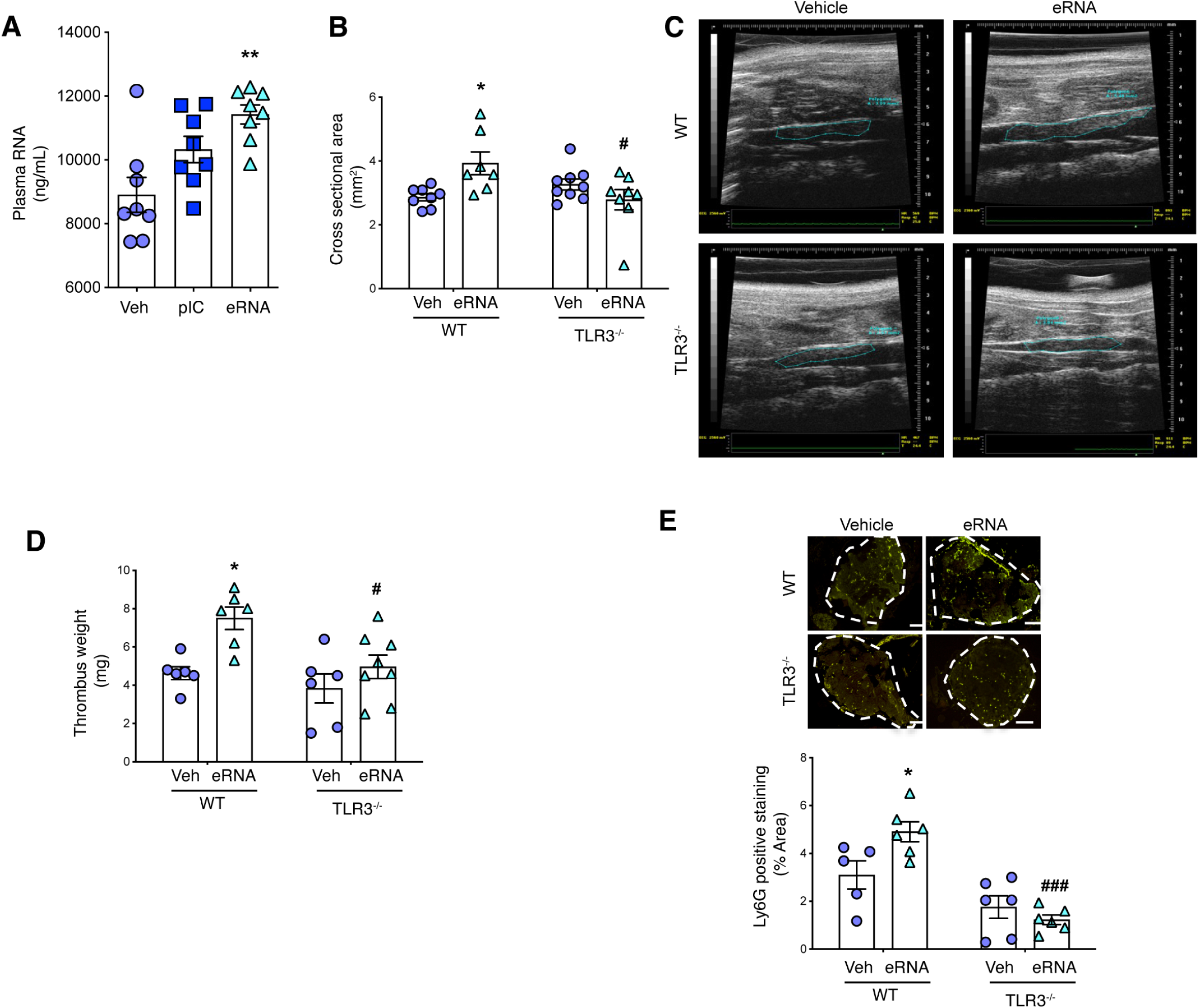
eRNA increase venous thrombosis. (A) eRNA levels in plasma are significantly increased following exogenous injection of RNA (in ng/mL) (N=8). (B) Injection of eRNA to WT mice increased thrombus cross sectional area measured using ultrasonography. eRNA increased thrombus size in WT mice. However, eRNA did not affect thrombus size in TLR3^-/-^ mice (N=7-9). (C) Representative ultrasound images of thrombosed veins in WT or TLR3^-/-^ mice treated or not with eRNA. (D) eRNA increased thrombus weight in WT but not in TLR3^-/-^ mice (N=6-7). (E) eRNA increased the infiltration of neutrophils in WT but not TLR3^-/-^ mice (magnification 10x; Bar=50 μm; N=5-6). *P<0.05 WT vehicle vs WT eRNA, #P<0.05 or ###P<0.001 WT eRNA vs TLR3^-/-^ eRNA.

The stasis model was also used to analyze the effect of poly(I:C), eRNA or RNase I treatments on thrombus parameters in WT animals. Analyses of Cartairs’ staining and ultrasonography showed that treatments did not affect thrombus size (Figure 4 A, B). Neutrophil infiltration was slightly enhanced by poly(I:C) and eRNA treatments; however, results did not reach statistical difference. (Figure 4C).

**Figure 4.**
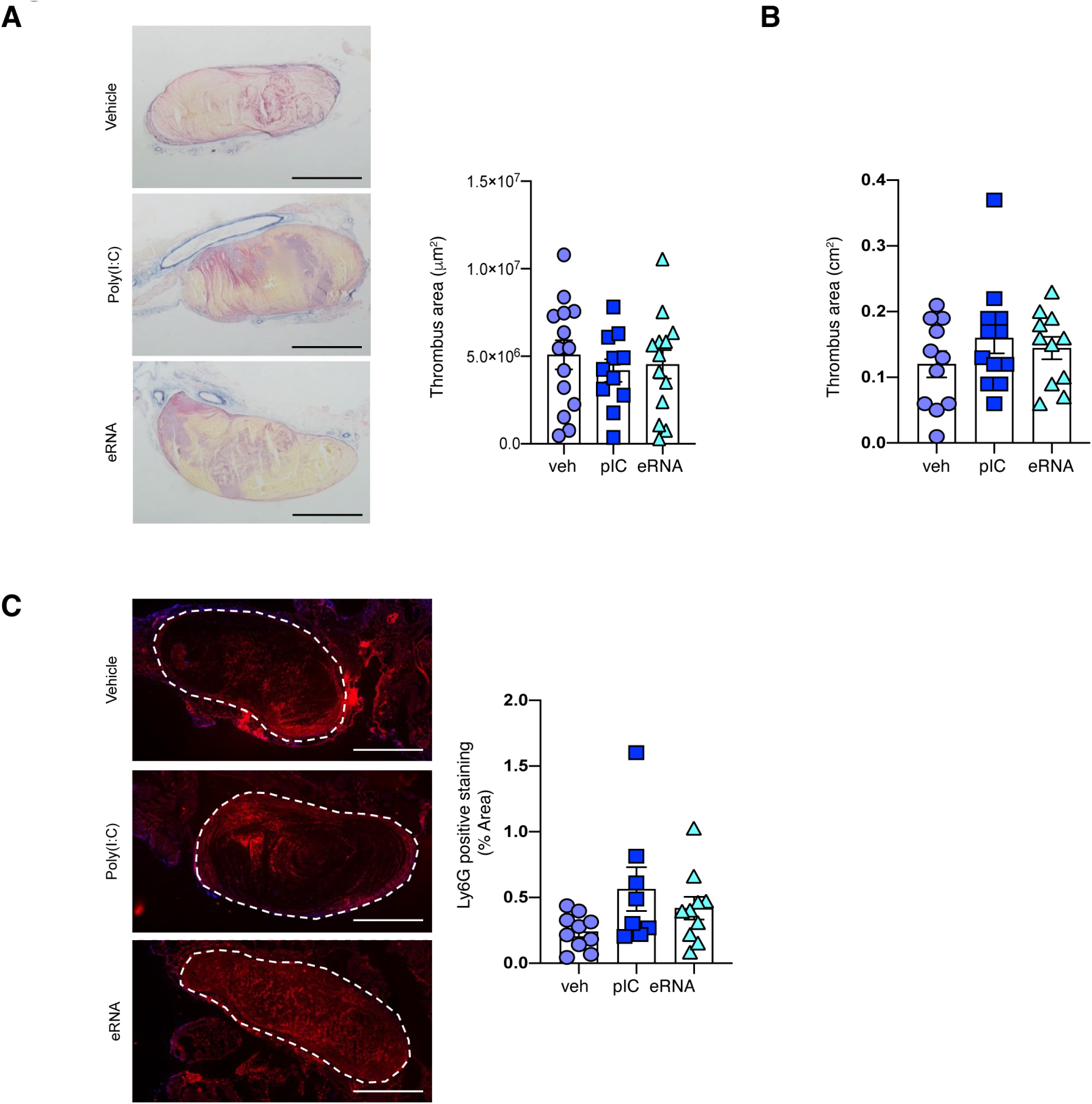
eRNA and poly(I:C) contribution to thrombosis in the stasis model. (A) Carstair’s staining representative images and quantifications 48h after thrombosis induction using the stasis model. Thrombus size determined by Carstair’s stained is similar in all experimental groups (magnification 10x Bar=50 μm N=11-14). (B) Thrombus area determined by ultrasonography showed equivalent results (N=11). (C) Ly6G staining representative images and quantifications in vehicle-, poly(I:C)- or eRNA-treated groups. Poly(I:C) and eRNA treatments were associated with a slight increase of neutrophil infiltration within thrombus, although not statistically different from vehicle-treated animals (magnification 20x Bar=25 μm; N=8-10).

Finally, we verified that RNase I, poly(I:C) or eRNA treatments did not affect DNA plasmatic concentrations in the FeCl_3_ or the stasis models of venous thrombosis as shown by DNA quantification in plasma of mice (Suppl Figure II).

### eRNA induce CXCL5 expression in endothelial cells

To identify the signals that coordinate the recruitment of neutrophils and favor their activation during venous thrombosis, HUVECs were treated with poly(I:C) in presence or absence of TLR3. HUVECs were transfected with a siRNA targeting TLR3 (siTLR3) or siRNA control (siCTR). TLR3 siRNA significantly reduced TLR3 expression in HUVECs. Interestingly, poly(I:C) treatment induced the expression of IL-8 and tissue factor (TF) in cells transfected with the siCTR. Downregulation of TLR3 in HUVEC significantly inhibited expression of these genes (Supplemental Figure III-A-C).

To determine if IL-8 produced by HUVECs after poly(I:C) incubation had a functional role on neutrophils, conditioned media from HUVECs transfected with siCTR or siTLR3 and treated with poly(I:C) were collected. The conditioned media was used to activate neutrophil migration in Boyden chambers. Neutrophil migration was increased following incubation with conditioned media from siCTR HUVEC stimulated with poly(I:C) compared to vehicle. Addition of an IL-8 receptor antagonist (SB225002) to the conditioned media significantly reduced neutrophil migration. Conditioned media from HUVECs transfected with siTLR3 with or without poly(I:C) or SB225002 did not alter neutrophil migration (Figure 5A).

**Figure 5.**
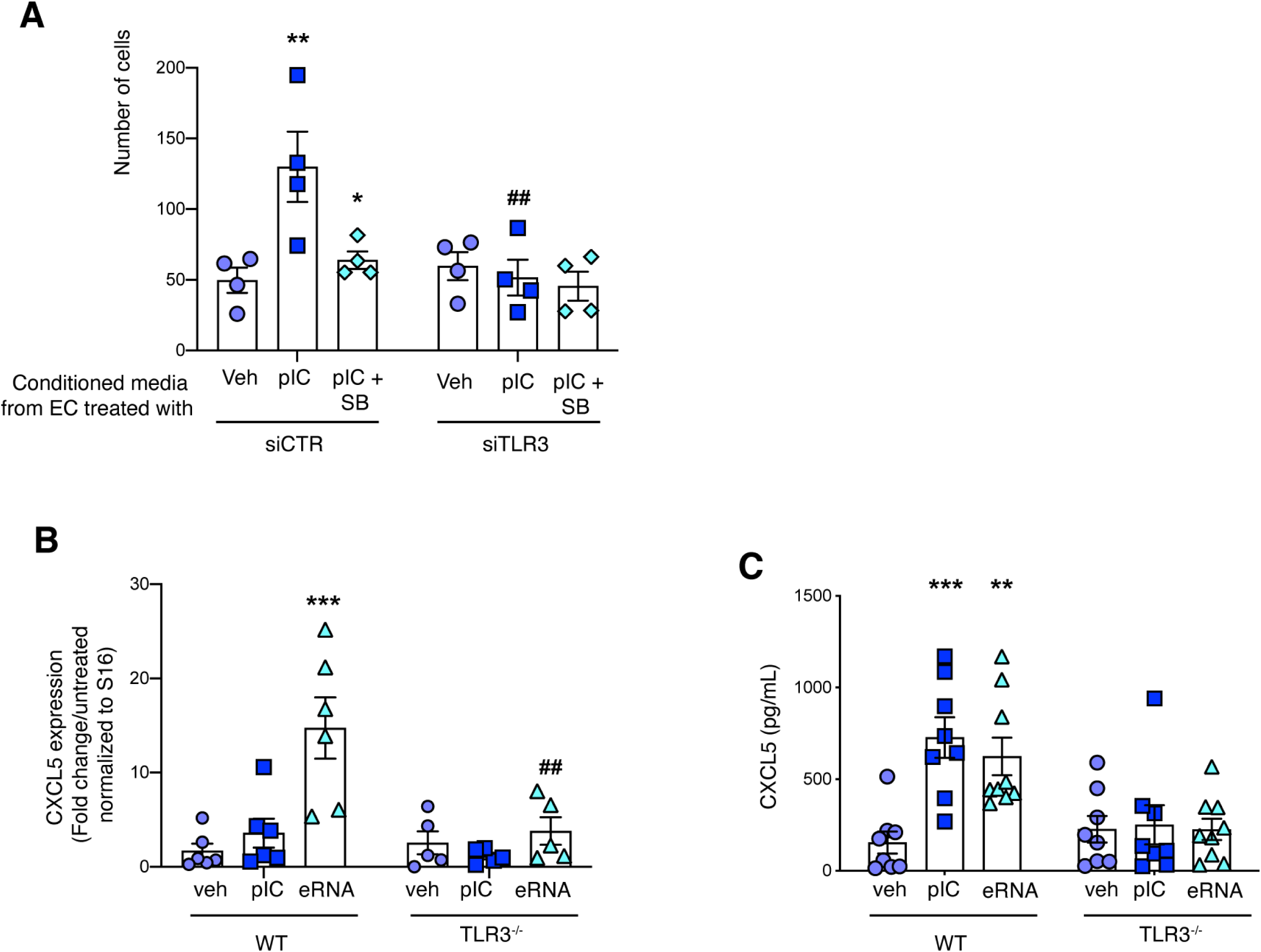
TLR3 activation promotes neutrophil migration and CXCL5 expression from endothelial cells. (A) Conditioned media from HUVECs treated with poly(I:C) induced neutrophil migration. This effect is dependent on CXCR2. On the contrary, conditioned media from HUVEC deficient for TLR3 did not modify the migratory capacity of neutrophils (N=4). (B) In murine endothelial cells, incubation with eRNA for 4h induced mRNA (N=5-6) and (C) protein (N=8-9) expression of CXCL5. The presence of TLR3 is required to promote CXCL5 expression. **P<0.01 vehicle vs treatment in HUVEC transfected with control siRNA; *P<0.05 siCTR poly(I:C) vs siCTR poly(I:C) + SB225002; ##P<0.01 siCTR vs siTLR3 transfected HUVEC treated with poly(I:C). ***P<0.001 WT vehicle vs WT eRNA; ##P<0.01 WT eRNA vs TLR3^-/-^ eRNA and *P<0.05 WT vehicle vs WT treated.

Mice lack a direct homologue of IL-8 but CXCL1/KC and CXCL5-6/LIX are considered functional homologues of IL-8 and participate to neutrophil recruitment^24^. Furthermore, CXCL1 and CXCL5 expression was found to be increased in mouse venous thrombi^4^. mRNA expression of CXCL1 was induced by poly(I:C) in WT but not TLR3^-/-^ endothelial cells; however, eRNA did not induced CXCL1 production from WT or TLR3^-/-^ endothelial cells (data not shown). Thus, we investigated if eRNA modified CXCL5 expression. Interestingly, we found that eRNA increased CXCL5 gene expression and protein secretion in WT endothelial cells. In endothelial cells deficient for TLR3, CXCL5 expression or secretion was not modified by eRNA treatment (Figure 5B-C).

### eRNA directly activate TLR3

Signaling through TLR3 classically leads to NFκB and IRF-3 activation and expression of type I IFN genes and pro-inflammatory cytokine. To directly demonstrate that eRNA activate TLR3, HEK293T cells were transfected with an expression plasmid for human TLR3 and incubated with either poly(I:C) or eRNA. TLR3 protein and mRNA expression were significantly increased in cells transfected with the TLR3 plasmid (Figures 6A, B). Western blot analysis showed that both poly(I:C) and eRNA significantly increased NFκB phosphorylation only in cells transfected with the TLR3 plasmid (Figure 6C). Additionally, in HEK293T overexpressing TLR3, NFκB activation induced by eRNA was associated with increased mRNA expression of interferon beta (IFNβ) (Figure 6D).

**Figure 6.**
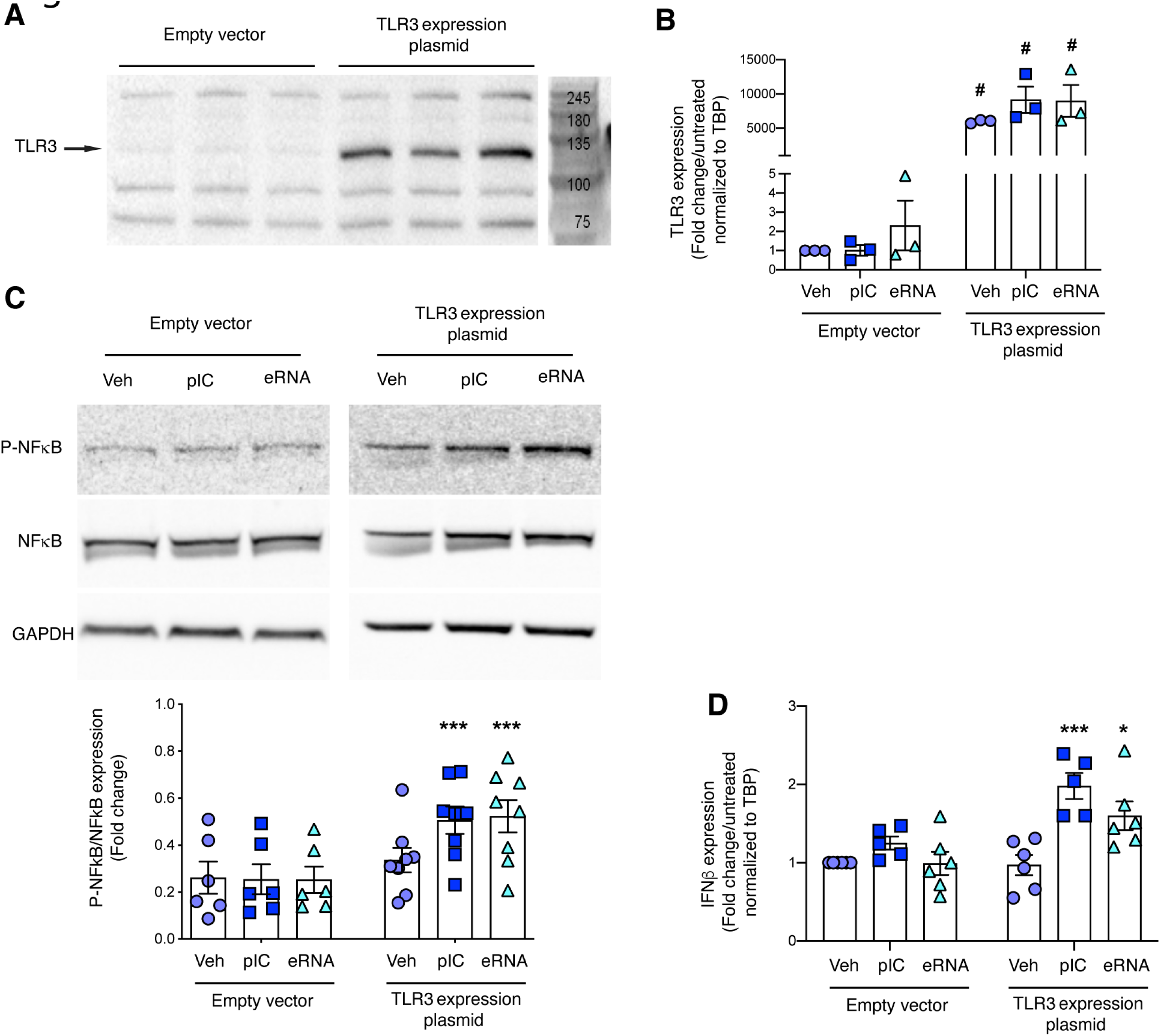
eRNA induce intracellular signaling pathways through TLR3 in HEK293T cells. (A) Transfection of HEK293T cells with an expression plasmid for human TLR3 significantly increased TLR3 protein expression compared to cells transfected with an empty vector (N=3). (B) TLR3 mRNA is also significantly expressed after 4h incubation with poly(I:C) and eRNA in HEK293T over-expressing TLR3 compared to vehicle. (C) 30 minutes of eRNA treatment of HEK293T cells over-expressing TLR3 significantly increased phosphorylation of NFκB. (D) This is associated with higher expression of TLR3 target gene IFNβ compared to unstimulated cells (4h treatment). *P<0.05 vehicle vs eRNA in HEK293T cells over-expressing TLR3, ***P<0.001 vehicle vs treatment in HEK293T cells over-expressing TLR3 and #P<0.05 empty vector vs TLR3 expression plasmid for equivalent treatment.

NFκB activation was also quantified in HUVECs transfected or not with a siRNA targeting TLR3 (Figure 7A). We found that poly(I:C) and eRNA treatments were associated with increased NFκB activation in HUVECs transfected with a control siRNA. Interestingly, eRNA no longer induced NFκB phosphorylation when TLR3 expression was downregulated by siRNA (Figure 7B).

**Figure 7.**
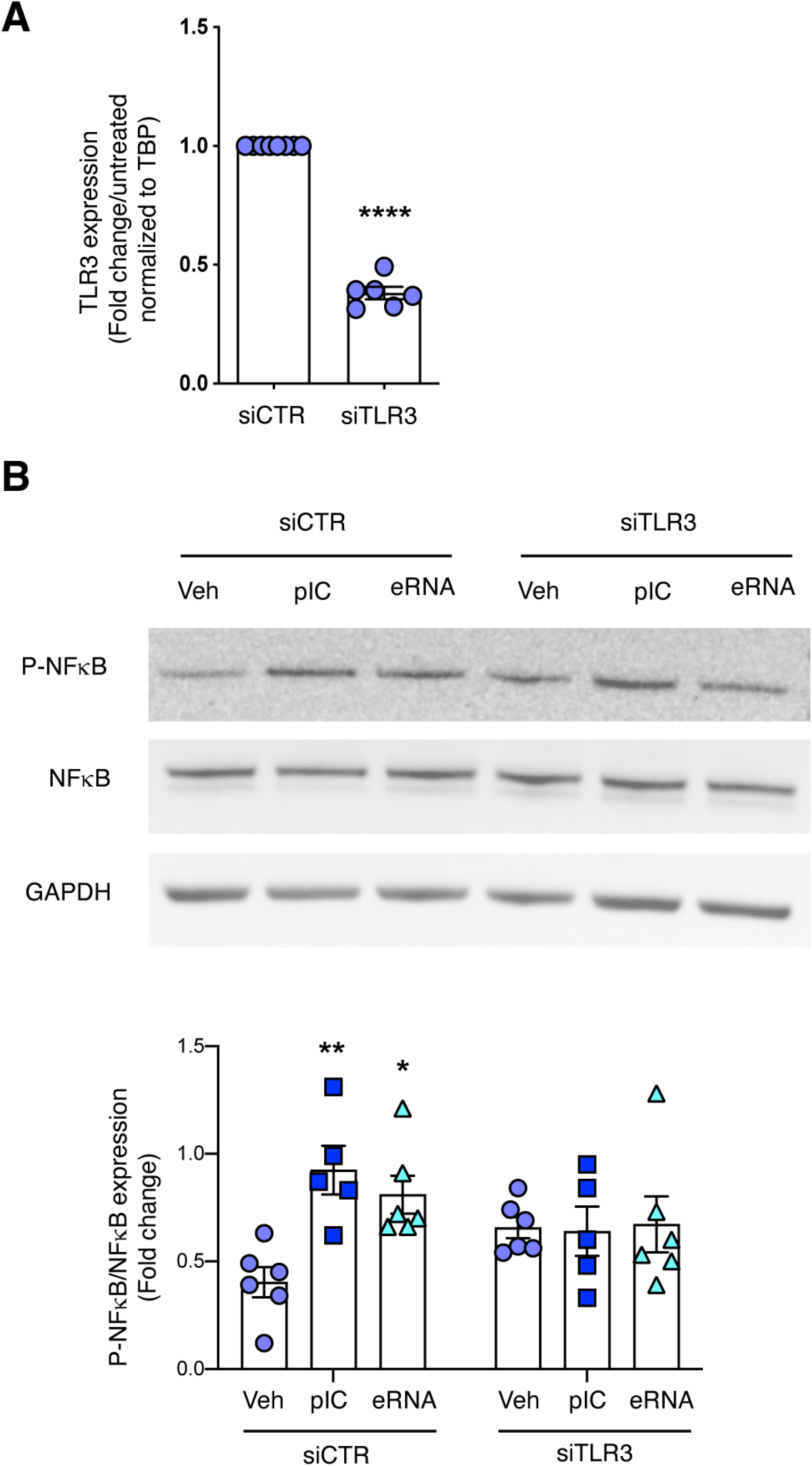
eRNA induce intracellular signaling pathways through TLR3 in HUVECs. (A) Transfection of HUVECs with an siRNA against TLR3 significantly reduced mRNA TLR3 expression (N=6-8). (B) Transfection of HUVEC with an siRNA against TLR3 blocked the eRNA-induced phosphorylation of NFκB after 45 minutes of treatment (N=5-6). **P<0.01 vehicle vs ploy(I:C) and *P<0.05 vehicle vs eRNA.

### eRNA induce coagulation and endothelial dysfunction through TLR3

To determine if eRNA-dependent activation of endothelial TLR3 modulates the coagulation cascade, plasma clot formation experiments were done on HUVECs. Re-calcified plasma was added to HUVECs following treatment with poly(I:C) or eRNA. Fibrin formation was measured at 405nm for 2h. Poly(I:C) and eRNA treatment significantly increased turbidity change and area under the curve (AUC) suggesting an amplification of fibrin network formation (Figure 8A-B). In addition, maximum absorbance was significantly increased by eRNA treatment (Figure 8B). We also found that TFPI expression was reduced in endothelial cells treated with eRNA compared to control (Figure 8C).

**Figure 8.**
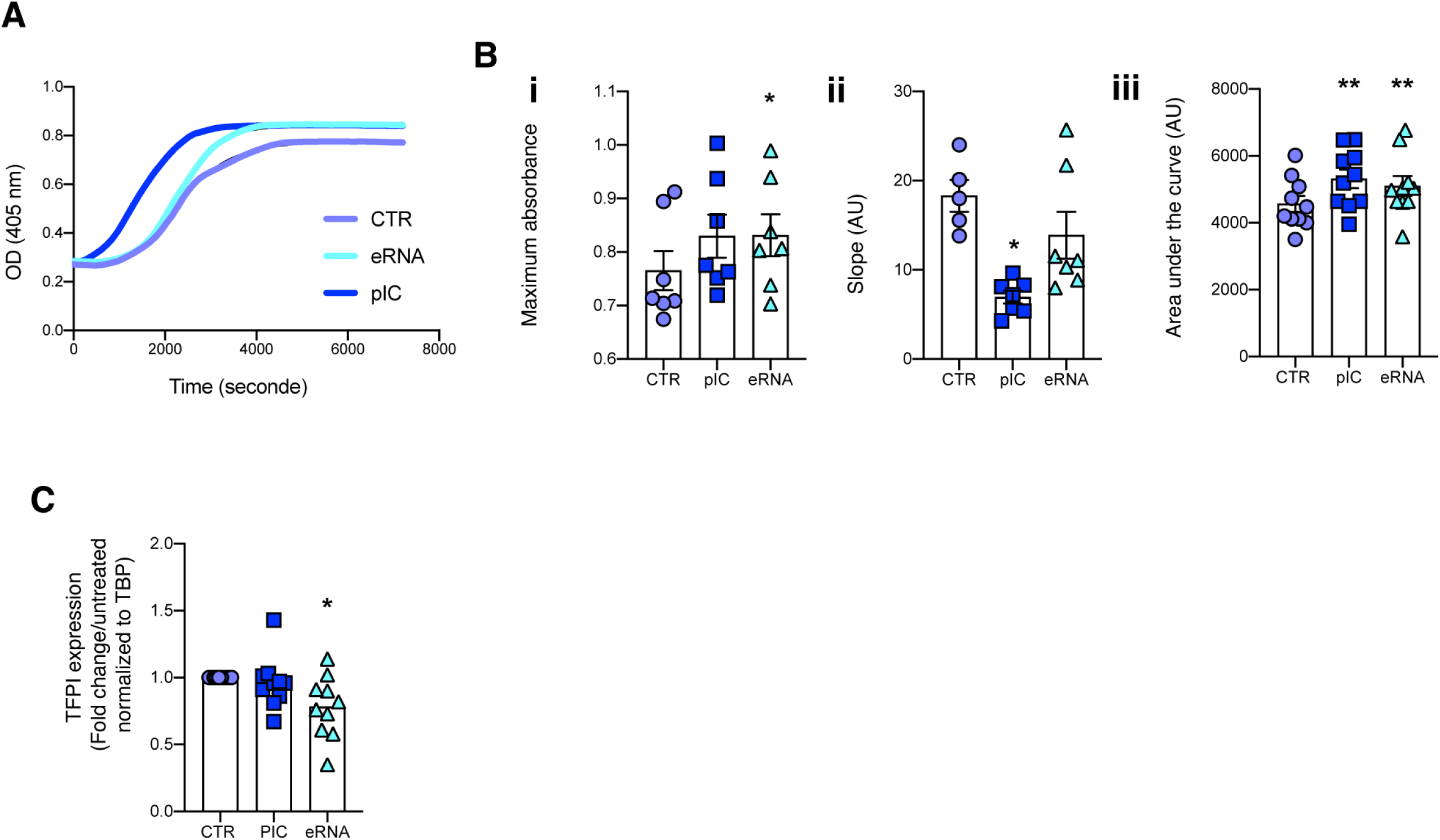
eRNA trigger coagulation. (A) Representative curve of clot formation assays on HUVEC monolayer. After incubation of HUVECs with eRNA or poly(I:C), clot formation was triggered in plasmas from healthy individuals by recalcification. Clot formation was monitored by turbidity. (B) eRNA significantly increased maximum absorbance (i, N=7) but the slope which represents the rate of fibrin formation was unchanged (ii, N=7). Interestingly, both eRNA and poly(I:C) significantly increased the area under the curve (AUC) (iii, N=10). (C) eRNA significantly reduced TFPI mRNA expression (N=8). *P<0.05 or **P<0.01 vehicle vs treatment.

## Discussion

In the present study, we demonstrate *in vivo* that eRNA promote venous thrombosis by activating TLR3. We showed *in vitro* that eRNA promote TLR3-dependent expression of a potent neutrophil chemoattractant cytokine, CXCL5.

eRNA derived from activated, stressed or injured cells under various pathological conditions such as hypoxia, infection, inflammation or tumor growth are detectable in body fluids. eRNA can be found within microvesicles or directly released from damaged cells. These free eRNA act as alarmins by inducing pro-coagulant and pro-inflammatory responses^25^. Compelling evidence suggest an important role of eRNA in cardiovascular diseases. Hence, eRNA are natural pro-coagulant co-factor, promote tissue and vessel regeneration, increase vascular permeability, induce pro-inflammatory mediator release from vascular and inflammatory cells and trigger cell recruitment^26^. In arterial thrombosis, eRNA act as primary inducers/cofactors of contact phase coagulation proteins, including factors XII and XI, as well as prekallikrein and high-molecular weight kininogen through direct polyanionic interactions. Since VTE is an immuno-inflammatory disease and that eRNA promote leukocyte recruitment and cytokine secretion, we sought to investigate its role during venous thrombosis.

Here we show that eRNA are released during venous thrombosis induced after vessel injury and participate in the pro-coagulant response. Interestingly, treatment with RNase I decreased neutrophil recruitment, which may explain the reduced thrombus size. Hence, neutrophils promote venous thrombosis through the formation of NETs, composed of DNA, histones and cytoplasmic proteins released from cytoplasmic granules^27^. These NETs contribute to venous thrombosis by promoting platelet aggregation and thrombin formation by providing a scaffold for procoagulant molecules including vWF, fibronectin, fibrinogen, FXII and TF and contribute to anticoagulant factor cleavage (TFPI and thrombomodulin)^28^. Here, we provide evidence for a beneficial role of RNase I regarding neutrophil recruitment and thrombus formation. In RNase I deficient mice, it was shown that circulating RNA were increased, which support activation of FXII and FXI. However, RNase I deficient mice did not show differences in clotting time compared to wild type animals. Nonetheless, it appears from this study that RNase I deficient animals might exert a susceptibility for thrombosis through FXII activation^29^. Here, we suggest that, beside direct effects on coagulation factors, RNA promote neutrophil activation and endothelial dysfunction leading to a pro-thrombotic phenotype.

Studies have suggested that TLR3 might have a role during thrombosis. Shibamiya et al. demonstrated that poly(I:C), a synthetic ligand of TLR3, upregulated TF and downregulated thrombomodulin expression in endothelial cells. *In vivo*, poly(I:C) elevated D-dimer levels, which suggested increased coagulation and fibrinolysis^12^. Other studies demonstrated that TLR3 activation in hypoxia models increased TF expression and activity, which suggest a pro-thrombotic role of TLR3. These effects were induced by poly(I:C) through TLR3^30,31^. Poly(I:C) also induced plasminogen activator inhibitor-1 (PAI-1) expression in human glomerular endothelial cells and this effect was abolished by the antimalarial agent chloroquine^32^. In addition, activation of TLR3 by poly(I:C) *in vivo* has been shown to impair endothelial function including vasodilation, ROS production and endothelial repair following injury^11^. More recently, TLR3 was found to contribute to the development of aortic valve stenosis^33^. In cancer, double-stranded RNA (dsRNA) released by disseminated tumor cells was detected by TLR3 causing endothelial SLIT2 induction, which promote cancer cell migration and intravasation toward endothelial cells^34^.

In contrast to the TLR3 detrimental effects mentioned above, studies have also elucidated beneficial roles of this receptor in pulmonary hypertension and idiopathic pulmonary fibrosis. Reduced expression of TLR3 contributes to endothelial apoptosis and pulmonary vascular remodeling leading to pulmonary hypertension^35^. Moreover, the TLR3 L412F polymorphism, which is characterized by defective TLR3 activation, is associated with aberrant fibroproliferation and accelerated progression of idiopathic pulmonary fibrosis^36^. Finally, in atherosclerosis, TLR3 has been shown to have protective effects in *in vivo* models of mechanical and hypercholesterolemic arterial injury^37^.

Here, we found that TLR3 activation induced by poly(I:C) increases venous thrombosis. This is partially explained by poly(I:C)-induced neutrophil accumulation in WT mice, compared to TLR3^-^/^-^ mice. Contrary to our result, Suresh et al. showed that TLR3 deficiency favors neutrophil recruitment, conferring a protective role during pulmonary infection^38^. This discrepancy could be explained by pathological differences and cellular type. Imhof et al. described that TLR3, with TLR4, exhibits early neutrophil arrival followed by late and weak monocyte recruitment. This study corroborates our data showing that TLR3 activation fosters neutrophil recruitment^39^. However, when thrombosis was induced by complete ligation of the IVC, treatments with poly(I:C) or eRNA were not associated with increased thrombus size or neutrophil infiltration compared to vehicle. It is possible that the lack of effect observed with the stasis model compared to the FeCl_3_ can be explained by the absence of blood flow associated with the ligation model. Hence, this might have masked or inhibited the effect of poly(I:C) and eRNA on thrombus formation^40^.

Reports suggest that short RNA sequences are sufficient to stimulate TLR3^9^. We showed that eRNA increase thrombus size and neutrophil recruitment through endothelial TLR3. Although we did not identify the source and the nature of the RNA in our *in vivo* model of venous thrombosis, our work corroborates that eRNA or self-RNA are indeed ligands of TLR3. Hence, our *in vitro* work demonstrates that eRNA-dependent NFκB phosphorylation require TLR3 in HEK293T cells and in HUVECs. Additional studies have suggested that eRNA are mediating their effects through TLR3^13,31,41–43^. For example, in myocardial infarction, eRNA released from damaged tissue were found to activate the TLR3-TRIF pathway^14^. Here, we found that eRNA induced classical target gene of TLR3, IFNβ, in HEK293T cells overexpressing TLR3.

Since human neutrophils do not express TLR3, we sought to investigate how eRNA may promote neutrophil recruitment and activation^43^. TLR3 and eRNA independently induce the release of pro-inflammatory cytokines in cardiovascular diseases^44^. Here, we found that poly(I:C) mediates IL-8 expression, which is a strong chemoattractant for neutrophils, from human endothelial cells. Downregulation of TLR3 expression using siRNA leads to a reduced IL-8 secretion from endothelial cells. Low level of IL-8 was associated with reduced neutrophil migration *in vitro*. CXCL1, CXCL5 and CXCL6 are considered as IL-8 homologues in mice. We found that secretion of CXCL5 from murine endothelial cells was induced by eRNA in a TLR3-dependent manner *in vitro*. Although we did not confirm the role of CXCL5 *in vivo*, one can speculate that CXCL5 is released from cytoplasmic granules of endothelial cells and platelets upon cell activation during thrombosis^24^. DeRoo et al. recently reported that the most upregulated genes in the vein wall following venous thrombosis in mice were inflammatory genes in comparison to sham operated animals. Importantly, Cxcl5 was one of the genes highly upregulated in VTE mice compared to sham animals^45^. Although expression of Cxcl5 was not attributed to the endothelial cell, it supports our results showing that CXCL5 production during VTE might be an important mechanism by which neutrophils are recruited within the thrombus.

Importantly, TLR3 and thrombin receptors, PAR1/2, were found to cooperate in the expression of pro-inflammatory and pro-thrombotic genes in endothelial cells^46^. Hence, we showed that eRNA-dependent activation of TLR3 is associated with increased coagulability of endothelial cells. Turbidity experiments demonstrated that treatment of HUVEC with poly(I:C) or eRNA enhanced fibrin formation in normal plasma. We showed that eRNA decreased TFPI expression^46^. Interestingly, it has been shown that NFκB activation was associated with reduced TFPI expression in HUVECs^47^. This mechanism may also be involved in our experimental conditions. Although we did not test it directly, one hypothesis is that eRNA might regulate TFPI expression through TLR3/NFκB activation. Further characterization of the mechanism would be required.

As it has been reported in previous studies, the role of TLR3 as a receptor for eRNA is still under debate and appears to be minor in mediating eRNA-cytokine responses in the context of arterial diseases^26^. However, in the venous system, eRNA induce mechanisms relevant for VTE through TLR3. We believe that these apparent discrepancies with our study may be due to different cell type and vascular beds but also the chemokines or cytokines studied. However, this would require further investigations.

In summary, we found that eRNA act through TLR3 to induce CXCL5 expression. We acknowledge that this mechanism would require more thorough investigation *in vivo*. Nonetheless, we clearly demonstrated that eRNA promote venous thrombosis through neutrophil recruitment in TLR3-dependent manner. Therefore, novel approaches counteracting or neutralizing damages induced by eRNA might represent interesting therapeutic options. Our study provides important evidence regarding the role of endogenous eRNA in the induction of pro-coagulant pathways over the course of venous thromboembolism.

## Author contributions

M. Najem and R. N. Rys performed research, analyzed data and wrote the manuscript. S. Laurance, F-R. Bertin, V. Gourdou-Latyszenok, L. Gourhant, L. Le Gall and R. Le Corre performed research and analyzed data. M. D. Blostein and F. Couturaud interpreted the data and wrote the manuscript. C. A. Lemarié conceived and supervised the study, performed research, analyzed data and wrote the manuscript.

## Acknowledgment

This study was supported by The Morris and Bella Fainman Family Foundation. C.A.L. was supported by Région Bretagne, Département du Finistère and Brest Métropole.

We would like to acknowledge and thank the animal care and flow cytometry (Hyperion) core facilities for their technical assistance.

## Disclosures

The authors have no conflict of interest.

**Supplemental Figure I. Thrombin-antithrombin in plasma of mice following thrombosis induction using the FeCl_3_ or the stasis models.** Thrombin-antithrombin (TAT) complexes were quantified in plasma of treated mice following thrombus induction with the FeCl_3_ (A) or the stasis (B) models. No statistical differences were found between all the experimental groups for each thrombosis models (N=5-8).

**Supplemental Figure II. Treatments do not modify DNA plasma levels in the FeCl_3_ or the stasis models.** (A) Circulating single-stranded DNA (ssDNA) was quantified in plasma of mice treated with poly(I:C) and eRNA in the FeCl_3_ model. Treatments did not change ssDNA plasma levels compared with controls. (B) Plasmatic ssDNA levels were also quantified in the stasis model in mice treated with poly(I:C), eRNA and RNAse I. No statistical differences were found between the experimental groups (N=8-11).

**Supplemental Figure III. TLR3 is required for gene expression in endothelial cells.** mRNA expression of TLR3 (A), IL-8 (B) and TF (C) in HUVEC transfected with an siRNA control or targeting TLR3. Poly(I:C) significantly enhanced expression of these genes in a TRL3-dependent manner (N=5). *P<0.05, **P<0.01 or ***P<0.001 vehicle vs treatment in HUVEC transfected with control siRNA, #P<0.05 or ##P<0.01 vehicle vs treatment in HUVEC transfected with TLR3 siRNA.

**Supplementary Figure IV. Uncropped version of Western Blot membranes for phospho-NFκB and NFκB in HEK293T cells transfected with an empty vector or with a human TLR3 expression plasmid.**

**Supplementary Figure V. Uncropped version of Western Blot membranes for phospho-NFκB and NFκB in HUVECs transfected with a control siRNA or a siRNA against TLR3.**

## Non-standard Abbreviations and Acronyms

DAMP: Damage-associated molecular pattern
DVT: Deep vein thrombosis
eRNA: Extracellular RNA
IVC: Inferior vena cava
NET: Neutrophil extracellular trap
PAMP: Pathogen-associated molecular pattern
TF: Tissue factor
TFPI: Tissue factor pathway inhibitor
TLR3: Toll-like receptor 3
VTE: Venous thromboembolism

## References

1. Fowkes, F. J., Price, J. F. & Fowkes, F. G. Incidence of diagnosed deep vein thrombosis in the general population: systematic review. Eur J Vasc Endovasc Surg 25, 1–5 (2003).

2. Kahn, S. R. et al. Determinants and time course of the postthrombotic syndrome after acute deep venous thrombosis. Ann Intern Med 149, 698–707 (2008).

3. Mehta, S. et al. Diagnostic Evaluation and Management of Chronic Thromboembolic Pulmonary Hypertension: A Clinical Practice Guideline. Canadian Respiratory Journal 17, 301–334 (2010).

4. von Bruhl, M. L. et al. Monocytes, neutrophils, and platelets cooperate to initiate and propagate venous thrombosis in mice in vivo. J Exp Med 209, 819–35 (2012).

5. Wagner, D. D. New links between inflammation and thrombosis. Arterioscler Thromb Vasc Biol 25, 1321–4 (2005).

6. Fuchs, T. A., Brill, A. & Wagner, D. D. Neutrophil extracellular trap (NET) impact on deep vein thrombosis. Arterioscler Thromb Vasc Biol 32, 1777–83 (2012).

7. Brill, A. et al. Neutrophil extracellular traps promote deep vein thrombosis in mice. J Thromb Haemost 10, 136–44 (2012).

8. Kariko, K., Ni, H., Capodici, J., Lamphier, M. & Weissman, D. mRNA is an endogenous ligand for Toll-like receptor 3. J Biol Chem 279, 12542–50 (2004).

9. Kleinman, M. E. et al. Sequence- and target-independent angiogenesis suppression by siRNA via TLR3. Nature 452, 591–7 (2008).

10. Alexopoulou, L., Holt, A. C., Medzhitov, R. & Flavell, R. A. Recognition of double-stranded RNA and activation of NF-kappaB by Toll-like receptor 3. Nature 413, 732–8 (2001).

11. Zimmer, S. et al. Activation of endothelial toll-like receptor 3 impairs endothelial function. Circ Res 108, 1358–66 (2011).

12. Shibamiya, A. et al. A key role for Toll-like receptor-3 in disrupting the hemostasis balance on endothelial cells. Blood 113, 714–22 (2009).

13. Cavassani, K. A. et al. TLR3 is an endogenous sensor of tissue necrosis during acute inflammatory events. J Exp Med 205, 2609–21 (2008).

14. Chen, C. et al. Role of extracellular RNA and TLR3-Trif signaling in myocardial ischemia-reperfusion injury. Journal of the American Heart Association 3, e000683 (2014).

15. Kannemeier, C. et al. Extracellular RNA constitutes a natural procoagulant cofactor in blood coagulation. Proc Natl Acad Sci U S A 104, 6388–93 (2007).

16. Fischer, S. et al. Extracellular RNA promotes leukocyte recruitment in the vascular system by mobilising proinflammatory cytokines. Thromb Haemost 108, (2012).

17. Simsekyilmaz, S. et al. Role of extracellular RNA in atherosclerotic plaque formation in mice. Circulation 129, 598–606 (2014).

18. Walberer, M. et al. RNase therapy assessed by magnetic resonance imaging reduces cerebral edema and infarction size in acute stroke. Current neurovascular research 6, 12–9 (2009).

19. Robins, R. S. et al. Vascular Gas6 contributes to thrombogenesis and promotes tissue factor up-regulation after vessel injury in mice. Blood 121, 692–9 (2013).

20. Aghourian, M. N., Lemarie, C. A. & Blostein, M. D. In vivo monitoring of venous thrombosis in mice. J Thromb Haemost 10, 447–52 (2012).

21. Aghourian, M. N., Lemarie, C. A., Bertin, F. R. & Blostein, M. D. Prostaglandin E synthase is upregulated by Gas6 during cancer-induced venous thrombosis. Blood 127, 769–777 (2016).

22. Bertin, F. R., Lemarie, C. A., Robins, R. S. & Blostein, M. D. Gas6 regulates thrombin-induced expression of VCAM-1 through FoxO-1 in endothelial cells. J Thromb Haemost (2015) doi:10.1111/jth.13156.

23. Pieters, M. et al. An international study on the feasibility of a standardized combined plasma clot turbidity and lysis assay: communication from the SSC of the ISTH. Journal of Thrombosis and Haemostasis 16, 1007–1012 (2018).

24. Hol, J., Wilhelmsen, L. & Haraldsen, G. The murine IL-8 homologues KC, MIP-2, and LIX are found in endothelial cytoplasmic granules but not in Weibel-Palade bodies. J Leukoc Biol 87, 501–8 (2010).

25. Preissner, K. T., Fischer, S. & Deindl, E. Extracellular RNA as a Versatile DAMP and Alarm Signal That Influences Leukocyte Recruitment in Inflammation and Infection. Frontiers in Cell and Developmental Biology 8, 1618 (2020).

26. Zernecke, A. & Preissner, K. T. Extracellular Ribonucleic Acids (RNA) Enter the Stage in Cardiovascular Disease. Circ Res 118, 469–79 (2016).

27. Rohrbach, A. S., Slade, D. J., Thompson, P. R. & Mowen, K. A. Activation of PAD4 in NET formation. Frontiers in immunology 3, 360 (2012).

28. Thålin, C., Hisada, Y., Lundström, S., Mackman, N. & Wallén, H. Neutrophil Extracellular Traps. Arteriosclerosis, Thrombosis, and Vascular Biology 39, 1724–1738 (2019).

29. Garnett, E. R. et al. Phenotype of ribonuclease 1 deficiency in mice. RNA 25, 921–934 (2019).

30. Biswas, I., Garg, I., Singh, B. & Khan, G. A. A key role of toll-like receptor 3 in tissue factor activation through extracellular signal regulated kinase 1/2 pathway in a murine hypoxia model. Blood Cells Mol Dis 49, 92–101 (2012).

31. Bhagat, S., Biswas, I., Ahmed, R. & Khan, G. A. Hypoxia induced up-regulation of tissue factor is mediated through extracellular RNA activated Toll-like receptor 3-activated protein 1 signalling. *Blood Cells*, Molecules, and Diseases 84, 102459 (2020).

32. Aizawa, T. et al. Chloroquine attenuates TLR3-mediated plasminogen activator inhibitor-1 expression in cultured human glomerular endothelial cells. Clin Exp Nephrol 23, 448–454 (2019).

33. Niepmann, S. T. et al. Toll-like receptor-3 contributes to the development of aortic valve stenosis. Basic Res Cardiol 118, 6 (2023).

34. Tavora, B. et al. Tumoural activation of TLR3-SLIT2 axis in endothelium drives metastasis. Nature 586, 299–304 (2020).

35. Farkas, D. et al. Toll-like Receptor 3 Is a Therapeutic Target for Pulmonary Hypertension. Am J Respir Crit Care Med 199, 199–210 (2019).

36. O’Dwyer, D. N., Armstrong, M. E., Kooblall, M. & Donnelly, S. C. Targeting defective Toll-like receptor-3 function and idiopathic pulmonary fibrosis. Expert Opinion on Therapeutic Targets 19, 507–514 (2015).

37. Cole, J. E. et al. Unexpected protective role for Toll-like receptor 3 in the arterial wall. PNAS 108, 2372–2377 (2011).

38. Suresh, M. V., et al. TLR3 absence confers increased survival with improved macrophage activity against pneumonia. JCI Insight 4, (2019).

39. Imhof, B. A., Jemelin, S. & Emre, Y. Toll-like receptors elicit different recruitment kinetics of monocytes and neutrophils in mouse acute inflammation. European Journal of Immunology 47, 1002–1008 (2017).

40. Diaz, J. A. et al. Choosing a Mouse Model of Venous Thrombosis. Arterioscler Thromb Vasc Biol 39, 311–318 (2019).

41. Bernard, J. J. et al. Ultraviolet radiation damages self noncoding RNA and is detected by TLR3. Nat Med (2012) doi:10.1038/nm.2861.

42. Green, N. M., Moody, K.-S., Debatis, M. & Marshak-Rothstein, A. Activation of Autoreactive B Cells by Endogenous TLR7 and TLR3 RNA Ligands*. Journal of Biological Chemistry 287, 39789–39799 (2012).

43. Jenne, C. N. et al. Neutrophils recruited to sites of infection protect from virus challenge by releasing neutrophil extracellular traps. Cell Host Microbe 13, 169–80 (2013).

44. Preissner, K. T. & Herwald, H. Extracellular nucleic acids in immunity and cardiovascular responses: between alert and disease. Thromb Haemost 117, 1272–1282 (2017).

45. DeRoo, E. et al. A vein wall cell atlas of murine venous thrombosis determined by single-cell RNA sequencing. Commun Biol 6, 1–13 (2023).

46. Subramaniam, S. et al. A thrombin-PAR1/2 feedback loop amplifies thromboinflammatory endothelial responses to the viral RNA analogue poly(I:C). Blood Advances 5, 2760–2774 (2021).

47. Chen, Y. et al. CRP regulates the expression and activity of tissue factor as well as tissue factor pathway inhibitor via NF-κB and ERK 1/2 MAPK pathway. FEBS Letters 583, 2811–2818 (2009).

